# Multi-environment phenotyping of *C. elegans* for robust evaluation of physical performance

**DOI:** 10.1101/2020.08.17.253583

**Authors:** Jennifer E. Hewitt, Ricardo Laranjeiro, Masoud Norouzi, Rebecca Ellwood, Adam Antebi, Nathaniel J. Szewczyk, Monica Driscoll, Siva A. Vanapalli

**Affiliations:** Department of Chemical Engineering, Texas Tech University, Lubbock, TX, USA; Department of Molecular Genetics of Ageing, Max Planck Institute for Biology of Ageing, and Cologne Excellence Cluster on Cellular Stress Responses in Aging-Associated Diseases (CECAD), University of Cologne, Cologne, Germany; Department of Molecular Biology and Biochemistry, Rutgers, The State University of New Jersey, Piscataway, NJ, USA; MRC/Arthritis Research UK Centre for Musculoskeletal Ageing Research, University of Nottingham, United Kingdom & National Institute for Health Research Nottingham Biomedical Research Centre, Derby, UK

## Abstract

Determining the physical performance of humans using several measures is essential to evaluating the severity of diseases, understanding the role of environmental factors, and developing therapeutic interventions. Development of analogous measures of physical performance in model organisms can help in identifying conserved signaling pathways and prioritizing drug candidates. In this study, we propose a multi-environment phenotyping (MEP) approach that generates a comprehensive set of measures indicative of physical performance in *C. elegans*. We challenge *C. elegans* in different mechanical environments of burrowing, swimming, and crawling, each of which places different physiological demands on the animals to generate locomotory forces. Implementation of the MEP approach is done using three established assays corresponding to each environment–a hydrogel-based burrowing assay, the CeleST swim assay, and the NemaFlex crawling strength assay. Using this approach, we study individuals and show that these three assays report on unique aspects of nematode physiology, as phenotypic measures obtained from different environments do not correlate with one another. Analysis of a subset of genes representative of oxidative stress, glucose metabolism, and fat metabolism show differential expression depending on the animal’s environment, suggesting that each environment evokes a response with distinct genetic requirements. To demonstrate the utility of the MEP platform, we evaluate the response of a muscular dystrophy model of *C. elegans dys-1* to drug interventions of prednisone, melatonin and serotonin. We find that prednisone, which is the current treatment standard for human Duchenne muscular dystrophy, confers benefits in all three assays. Furthermore, while the tested compounds improve the physical performance of *dys-1*, these compounds are not able to fully restore the measures to wild-type levels, suggesting the need for discovery efforts to identify more efficacious compounds that could be aided using the MEP platform. In summary, the MEP platform’s ability to robustly define *C. elegans* locomotory phenotypes demonstrates the utility of the MEP approach toward identification of candidates for therapeutic intervention, especially in disease models in which the neuromuscular performance is impaired.

## INTRODUCTION

Physical performance or fitness in humans is formally assessed using health-related or skill-related measures [1]. Health-related measures include cardiorespiratory endurance, muscle endurance, muscle strength, body composition, and flexibility. Skill-related measures include agility, balance, coordination, power, speed and reaction time. Assessment of human physical performance utilizing such measures is central to evaluating severity of diseases, especially diseases in which the neuromuscular system is involved [2–4]. Likewise, declines in physical performance have been shown to correlate with the risk of disability, hospitalization rates, and mortality in the elderly [5–8].

Linking the molecular, cellular, and tissue-level mechanisms to human physical performance is best achieved in model organisms as they allow identification of conserved signaling pathways and therapeutic interventions. The nematode *Caenorhabditis elegans* is a popular model organism that features relatively simple and well-understood biology, rapid lifecycle, and ease of culture [9]. Importantly, genetic conservation supports the relevance of the *C. elegans* model for studying gene activities and pathways relevant to a variety of human diseases [10–16] including Duchenne muscular dystrophy [17], neurodegenerative diseases [18–20], and sarcopenia [21]. Moreover, *C. elegans* has a conserved muscular architecture comprising contractile elements and attachments similar to higher animals. Thus, development of approaches to assess the physical performance of *C. elegans* using measures that are analogous to human measures may help in translating discoveries made in *C. elegans* to human health.

Assessment of physical performance in the nematode *C. elegans* usually involves monitoring the locomotion of crawling *C. elegans* on agar plates or thrashing in liquid [22, 23], although more recently, assays have also been developed to characterize the burrowing ability of *C. elegans* [24, 25]. Technical advances in computer vision and data-driven approaches have led to increasing the depth of phenotypic information from crawling and swim assays by generating multidimensional readouts [26, 27]. Despite these significant advances, current approaches to characterize *C. elegans* physical performance commonly rely on investigating animal locomotory response in a singular mechanical environment, limiting the extent of physiological and clinically relevant measures that can be obtained.

Depending on the mechanical resistance of the environment, *C. elegans* can adjust its gait and body mechanics [28]. On agar plates, the surface tension of the thin liquid layer at the agar surface constrains the animal motion to two dimensions and also aids in propulsive thrust [29]. In liquid, the nematode body experiences rotational slip and therefore to effectively swim, the internal forces generated within the animal need to balance the external hydrodynamic forces [30]. In gel environments, the animal burrows by executing three-dimensional maneuvers, making the neuromuscular actuation different from that on two-dimensional substrates [30, 31]. Finally, studies have shown that in microfluidic pillar environments, animals can adjust body gait, speed, and force generation depending on the geometry and mechanical resistance of the pillar arena [32–34]. Thus, opportunities exist to expose *C. elegans* to different mechanical environments and extract information analogous to human health-related or skill-related measures.

In this study, we establish a multi-environment phenotyping (MEP) platform that evaluates the physical performance of *C. elegans* in three distinct mechanical environments. Our approach combines three assays of physical performance — NemaFlex [34], CeleST [35], and a hydrogel-based burrowing assay [25] — to assess animal physical capacities in environments that elicit locomotory modes of crawling, swimming, and burrowing, respectively. By assaying the same individual animal in each environment, we show that each assay provides distinct information on animal capabilities, and that crawling, swimming, and burrowing evoke distinct responses associated with changes in expression of different genes. To demonstrate the applicability of the MEP platform for evaluating *C. elegans* disease models and therapeutic interventions, we assess the physiological health of *dys-1* mutants, the *C. elegans* model for Duchenne muscular dystrophy. We show that while the tested compounds improve *dys-1* health, the treatments are not able to fully restore wild-type behaviors, underscoring the need for further discovery efforts for compounds that counter defects in all the measures reported in the MEP platform.

## RESULTS

### Overview of the multi-environment phenotyping approach

The multi-environment phenotyping (MEP) platform consists of three assays conducted in different mechanical environments that evoke one of three forms of locomotion from *C. elegans*: burrowing, swimming, or crawling. We extracted descriptive measures from each of the three environments (Figure 1).

**Fig. 1.**
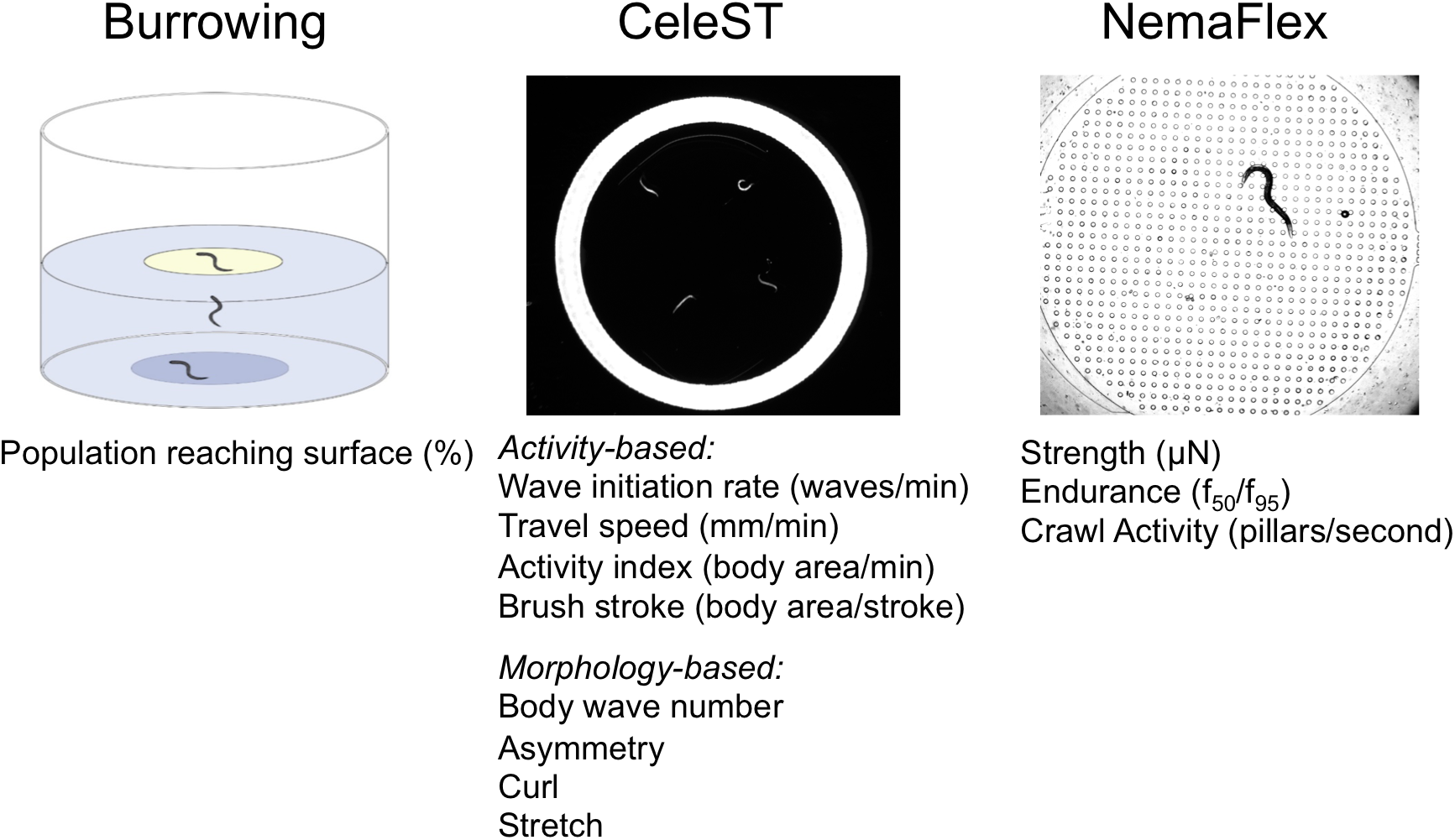
Overview of the multi-environment phenotyping (MEP) platform for evaluation of the physical performance of *C. elegans*. The MEP platform consists of a hydrogel-based burrowing assay, the CeleST swim assay, and the NemaFlex assay. From these three platforms, animal populations are assessed for their capacity to burrow in 3D, swim, and crawl, respectively. A total of twelve different measures are extracted from the three distinct mechanical environments.

In the hydrogel burrowing assay [25] we load animals into the base of a well and add a layer of Pluronic F-127 solution, which is a transparent and biocompatible hydrogel that transitions from liquid to solid under temperature upshifts. Pluronic solution is maintained at 14°C before transfer to the well plate, which allows the solution to stay liquid until it equilibrates to room temperature and gels to trap the nematodes. A chemoattractant is then added to the surface, encouraging the animals to burrow to the surface of the gel. Individuals can be scored based on the time they take to burrow to the surface, while with populations, the number of animals reaching the surface is counted at regular time intervals.

The second physical environment we tested is liquid, in which swim motion is assayed using the *<u>C.</u> <u>ele</u>gans* <u>S</u>wim <u>T</u>est (CeleST), analyzed with software that evaluates 8 measures related to body morphology and activity [35, 36]. The four morphological measures include (i) body wave number – a measure of waviness of the body posture, (ii) asymmetry – the degree to which the animal bends more toward one side or the other, (iii) stretch – a measure of how deep or flat the body bends are, and (iv) curling – percentage of time that an animal spends overlapping with itself. The four activity-related measures include (i) wave initiation rate – the number of body waves initiated by the head or tail per minute, (ii) travel speed – the distance travelled during a defined time, (iii) brush stroke – area painted by the animal body in a complete stroke, and (iv) swim activity index – brush stroke normalized by the time taken to perform two strokes.

The third environment features pillars that are deformed while the animal crawls in a microfluidic chamber, enabling calculation of muscle forces [34]. This NemaFlex assay involves imaging in a microfluidic device in which the nematode adopts a crawling gait due to the presence of a resistive array of flexible micropillars [34, 37]. From these images, pillars with maximum deflections are extracted, and these deflections are translated to maximal forces. Muscle strength is defined as the 95^th^ percentile of the maximal forces or *f*_*95*_. In addition to measuring the muscle strength from the NemaFlex assay, we also extract measures of muscular endurance and crawling activity. We define muscular endurance as the ratio *f*_*50*_/*f*_*95*_, where *f*_*50*_ represents 50^th^ percentile of the maximal forces. Animals that exert a high force as a one-time event would have a low ratio while those exerting large forces continuously would have a high ratio. Thus, the ratio *f*_*50*_/*f*_*95*_ is taken as a measure of muscular endurance. The crawling activity is calculated from the number of microfluidic pillars that the animal interacts with per unit time.

### An individual’s physical performance is distinct in different mechanical environments

Given that the burrowing, CeleST, and NemaFlex assays expose *C. elegans* to different mechanical environments, we wondered whether an individual that performs well in one environment would also perform better in a different environment. For example, if an animal shows good swim performance based on CeleST measures, the question is whether the same individual would show improved muscle strength as reported by the NemaFlex assay. Likewise, if an animal is efficient at burrowing and reaching the gel surface quickly, will this individual also display better swim performance? In addition, we sought to determine whether measures of physical performance obtained from different environments correlated with one another.

To address how different the physical performance of an individual is in each of our distinct mechanical environments, we phenotyped the same animal using burrowing, CeleST, and NemaFlex assays. We loaded wild-type animals into well plates with the pluronic gel, allowed them to burrow toward the surface, and measured the burrowing time for each individual. As each animal reached the gel surface, we immediately transferred it to a seeded NGM plate. The individuals were allowed to recover for 4 hours to allay any potential burrowing fatigue. We then transferred each individual into a liquid drop briefly for the CeleST assay while maintaining the identity of each animal from the burrowing assay. Finally, each of the indexed individuals was loaded into the NemaFlex device chambers for characterization of crawling and strength measures.

To assess inter- and intra-environment dependence, we determined the Pearson correlation coefficients for each combination of score metrics (Figure 2A). The correlation values between all variables are represented as the intensity value of the heat map. We find some strong correlations among the CeleST measures extracted from the swimming environment. Morphology-related measures of body wave number, asymmetry, and stretch all positively correlate with one another, while activity-based measures of wave initiation rate, travel speed, brush stroke, and swim activity also positively correlate with one another. Variables between the two morphology- and activity-related categories are negatively correlated with one another. Since it has previously been reported that the values of the morphological measures tend to increase with age and the activity-based measures tend to decrease in value with age [35], wild-type aging is at least one context in which these measures are correlated in a similar manner at a population level. Crawling-associated measures within the NemaFlex environment correlate weakly with one another.

**Fig. 2.**
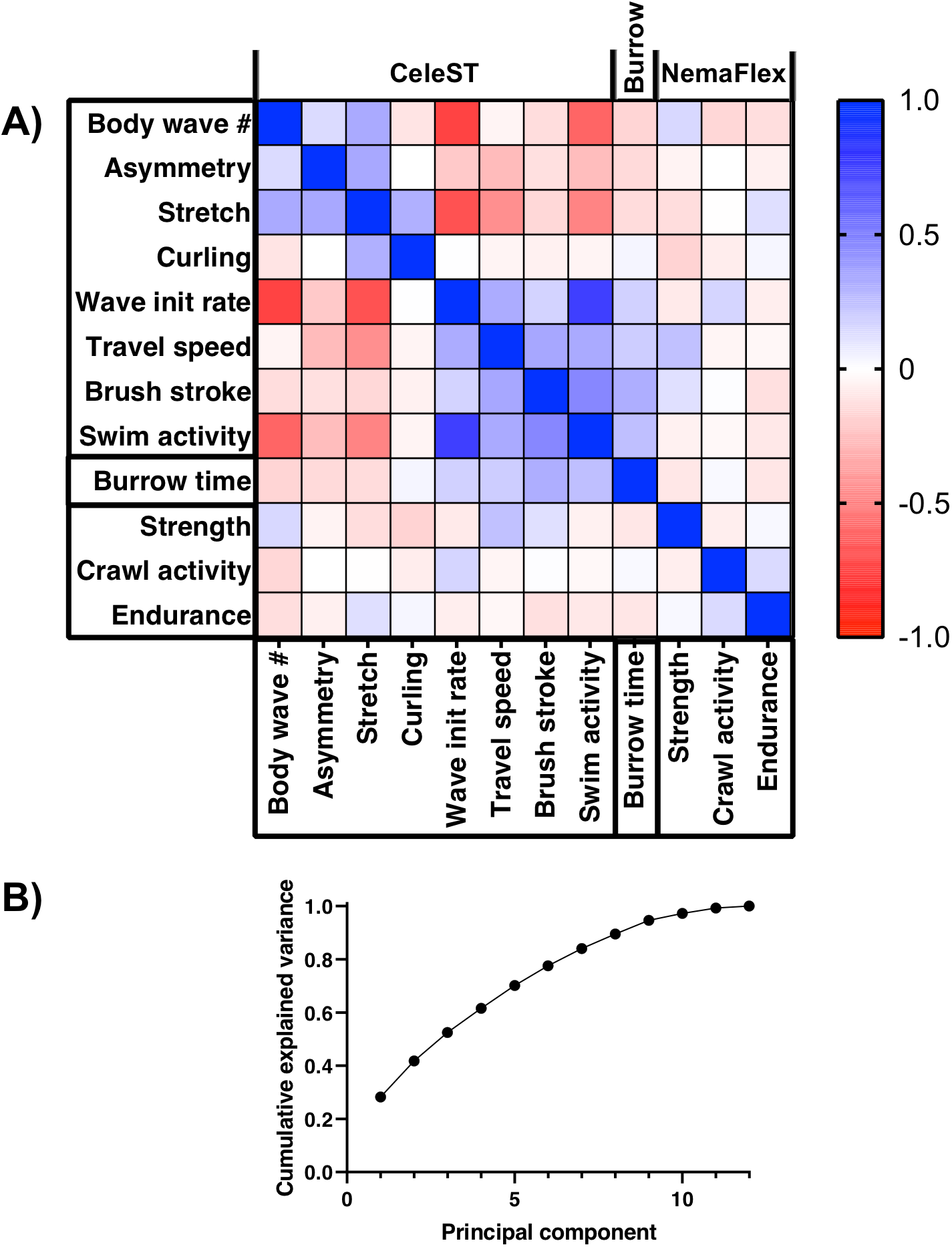
Measures extracted from the mechanically distinct swimming, crawling, and burrowing environments are not strongly correlated with one another. (A) A heat map shows the correlations among the different parameters extracted from the three different environments used to phenotype the same individual animals. Among all measures, strong correlations exist between only subsets of the CeleST swim measures, and not between the swimming, crawling, and burrowing measures. Intensity values in the heat map show the correlation coefficient obtained from a linear fit of the individual data for each variable plotted against one another. N=56 individual animals tested in all three environments. (B) A principle components analysis based on the correlation matrix does not reduce the dimensionality of the data, as seven components are necessary to explain 80% of the variation. This means that the data do not form any meaningful smaller sub-dimension in the original 12-dimensional space and therefore report primarily on distinct rather than inter-dependent measures.

Importantly, measures from different environments have correlation coefficients hovering around zero—indicating the different environments in fact reveal different and non-overlapping capabilities. Furthermore, principal component analysis (PCA) based on the correlation matrix of all variables shows that each principal component can explain only a small amount of the variance, with the first component accounting for less than 30% of the variance (Figure 2B). In fact, seven components are required to explain at least 80% of the variance of the dataset, indicating that no useful reduction of the dimensionality of the dataset is obtained via a PCA. Our data support the idea that the various measures reported from each of the environments reflect distinct indicators of an animal’s physiological health. The results also show that an individual performing better in one environment may not perform well in a different environment, suggesting that the MEP platform can comprehensively inform on *C. elegans* physiology.

### Crawling, swimming, and burrowing environments each elicit a unique transcriptional response

Our observations prompted us to ask whether animals exposed to each environment show differences at the gene expression level. We previously reported that placing *C. elegans* in a swimming environment as compared to a crawling environment elicits a specific gene expression response across a number of different functional clusters of genes, including those affecting oxidative stress response, glucose metabolism, and fat metabolism [38]. Documenting different transcriptional signatures in swimming versus crawling environments would support the value of assaying animals in different environments, as this multi-behavioral test might expose defects or improvements apparent in only a subset of environments.

To test for transcriptional profile intersections, we extended our qPCR data that compared changes in gene expression after 90 minutes of swimming to those in a control group of crawling worms [38] by evaluating selected expression in animals after 90 minutes of burrowing relative to a crawling control. Data for both swimming and burrowing animals are shown as the log_2_ fold change relative to a crawling group that was left on unseeded NGM plates during the 90-minute swim or burrowing period (Figure 3).

**Fig. 3.**
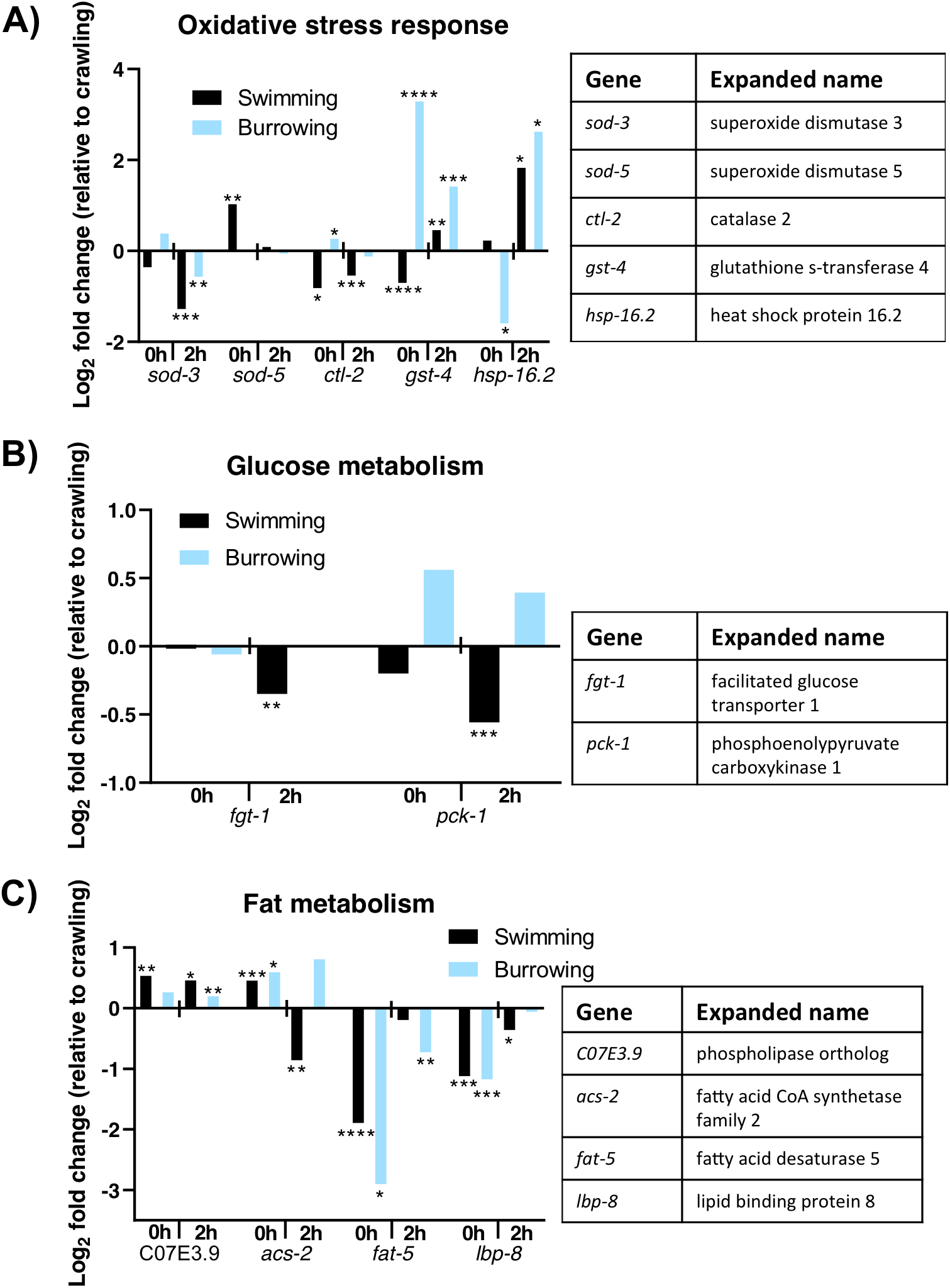
Swimming, burrowing, and crawling each elicit different gene expression responses. Animals that swim or burrow for 90 minutes differentially express genes involved in (A) oxidative stress response, (B) glucose metabolism, and (C) fat metabolism. Changes in expression are compared to a control group of animals that crawled on unseeded NGM plates for the 90-minute period. 0h is immediately after the 90-minute period; 2h is 2 hours after cessation of the activity trial period. With the permission of the authors, gene expression data from animals after 90 minutes of swimming is reanalyzed from published data in Ref. 38. For each of burrowing and crawling control populations, ~30 animals were collected, and experiments were done in triplicate. Significance was assessed with a 2-sample t-test. *, P≤0.05; **, P≤0.01; ***, P≤0.001; ****, P≤0.0001.

First, in animals that had burrowed for 90 minutes, we assessed five of the previously reported oxidative stress response genes that respond to swimming (*sod-3*, *sod-5*, *ctl-2*, *gst-4*, and *hsp-16.2)* (Figure 3A). Immediately after the burrowing or swimming at the 0h time point, there were some notable differences between the swimming and burrowing animals. While the superoxide dismutase 3 (*sod-3*) gene was not differentially regulated in burrowing vs. swimming animals, *sod-5* was significantly upregulated in animals that swam but was not changed in animals that burrowed. Catalase (*ctl-2)* and *gst-4* were significantly upregulated in burrowing animals but downregulated in swimming animals, while *hsp-16.2* was not changed in animals that swam but was largely downregulated in animals that burrowed. These results indicate that burrowing animals have an immediate transcriptional oxidative stress response distinct from that of crawling or swimming animals.

At the 2-hour time point, oxidative stress-related expression was fairly similar between animals that swam or burrowed. The time point 2 hours after challenge cessation may begin to reflect longer term adaptive changes. Notably, 2 hours after swim cessation, animals that swam downregulated the expression of glucose metabolism genes *fgt-1* and *pck-1,* which are involved in glucose transport and gluconeogenesis, respectively. Neither *fgt-1* nor *pck-1* was downregulated by burrowing (Figure 3B). Finally, the fat metabolism genes assessed here (C07E3.9, *acs-2*, *fat-5*, and *lbp-8*) showed a similar overall pattern in expression changes between burrowing and swimming animals; however, swimming and burrowing animals showed remarkably different expression patterns than their crawling counterparts (Figure 3C).

Taken together, analysis of a subset of genes with roles in the oxidative stress response, glucose metabolism, and fat metabolism supports the idea that the environments/experiences of burrowing, swimming, and crawling place distinct demands on animal physiology. This analysis, along with the individual-level physical performance phenotyping data from Figure 2, supports potential utilization of MEP to more comprehensively assess an animal’s physiological health.

### Application of MEP platform to *dys-1* mutants and their response to drug interventions

We have shown that the MEP platform is suitable for assessing the physical performance of *C. elegans* by exposing the animals to different mechanical environments. To demonstrate the utility of our approach, we utilized the MEP framework to assess a *C. elegans* disease model for Duchenne muscular dystrophy (DMD), in which the nematode physical performance is greatly impaired and this decline in physical performance is of clinical relevance. For this purpose, we chose *dys-1(eg33)* mutants that are defective in the *C. elegans* dystrophin gene. We were also curious to know which of the measures of physical performance would be improved when *dys-1* animals were treated with drugs that have been previously tested in worm and mouse models of DMD. We chose prednisone and melatonin as drug interventions since prednisone is the standard of care for DMD in humans [39], and melatonin has been tested in humans with some beneficial effects on disease pathology [40]. We also included serotonin, as serotonin was shown to be capable of preventing muscle damage in a *C. elegans* DMD model [41], although serotonin alone was not found efficacious in a follow-up study in a mouse DMD model [42].

#### Crawling environment in NemaFlex devices

Using NemaFlex, we previously reported that *dys-1(eg33)* mutants are significantly weaker than their wild-type counterparts on the third day of adulthood [37]. Here, using a microfluidic device optimized for younger adult animals, we show that *dys-1* mutants are significantly weaker than wild-type animals on the second day of adulthood (Figure 4A). The ability to detect this weakness at an early time point is relevant in that patients with DMD exhibit muscle weakness early in life and intervention testing in the worm model can now be focused on earlier outcomes.

**Fig. 4.**
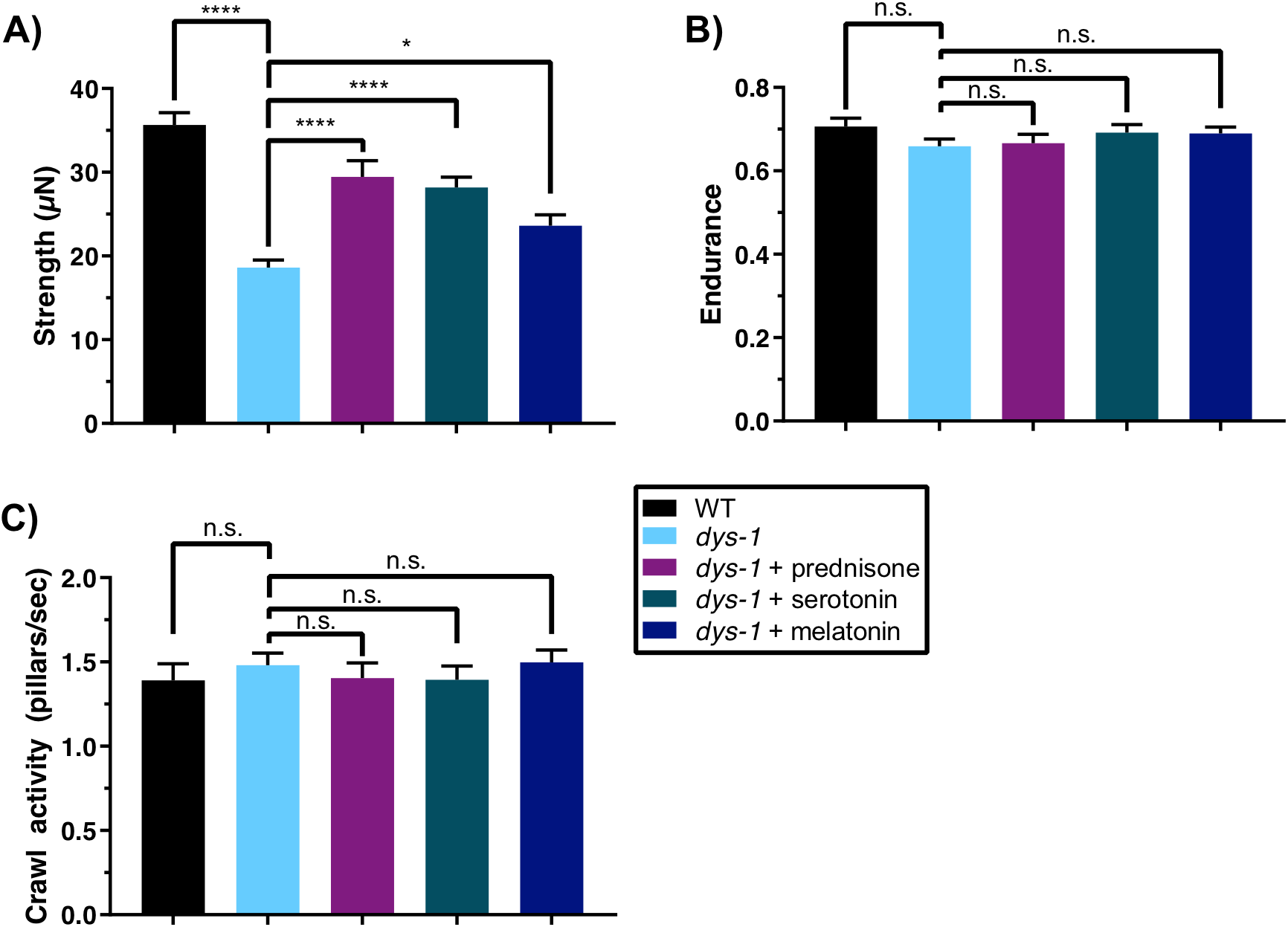
NemaFlex crawling environment: dystrophin mutants are deficient in their strength, which is improved by prednisone, serotonin, and melatonin. (A) We measured strength of animals crawling in a pillar matrix push against pillars in the NemaFlex microfluidics device. *dys-1(eg33)* mutants are deficient in their strength compared to wild-type animals. All three compounds tested improve animal strength, with prednisone yielding the largest improvement. (B) Endurance of *dys-1(eg33)* mutants is not significantly different from that of wild type and also remains unaffected under treatments. (C) The crawling activity, which is defined as the number of pillars interacted with per second, of *dys-1(eg33)* is also not different from wild type and does not change under any treatments. WT: N=28; *dys-1*: N=37; *dys-1*+prednisone: N=33; *dys-1*+serotonin: N=35; *dys-1*+melatonin: N=37. Significance between WT and *dys-1* controls was assessed with a 2-sample t-test; significance between *dys-1* and *dys-1*+treatments was assessed with one-way ANOVA with Dunnett’s post-hoc test. n.s., not significant; *, P≤0.05; ****, P≤0.0001.

With an interest in measuring the efficacy of clinical interventions aimed at alleviating the *dys-1* pathophysiology, we found that treating *dys-1* mutants with prednisone and melatonin induces significant improvements in muscle strength over the control (Figure 4A). Serotonin, which was previously reported to reduce the number of damaged muscle cells in the sensitized *dys-1;hlh-1* model [41], also improves muscle strength of the *dys-1* mutant. While all three compounds improve strength of the *dys-1* mutant, none are able to restore strength back to wild-type levels in the strength assay, indicating these treatments do not fully reverse the muscular defects of the *dys-1* mutant. In addition, we find that measures of muscle endurance and crawling activity are not significantly different in the *dys-1* mutant and also do not change under treatment with any of the compounds (Figure 4B-C). Thus, treatments that confer positive effects in humans also confer positive effects in *C. elegans*, but differences are not revealed in all the measures from NemaFlex assay.

#### CeleST swimming environment

*dys-1* mutants are deficient in their thrash rate in liquid, a single-parameter readout of swimming ability [37]. We were therefore interested in assessing whether the various activity- and morphology-related measures of CeleST were also deficient in swimming *dys-1* animals. We found a striking difference between *dys-1* mutants and wild-type animals in all four activity-related measures, such that brush stroke, activity index, travel speed, and wave initiation rate are each a small fraction of baseline values for *dys-1* as compared with control animals (Figure 5A-D). Prednisone, serotonin, and melatonin, each of which improves muscle strength of *dys-1* mutants, also confer modest but significant improvements in each of these four activity-related swim measures. Additionally, while *dys-1* mutants do not appear to spend more time in a curled morphology compared with wild-type animals (Figure 5E), they do have significantly higher asymmetry, stretch, and body wave number (Figure 5F-H). Prednisone, serotonin, and melatonin are able to restore body wave number closer to wild-type levels but have no significant effect on the abnormally high asymmetry of *dys-1* animals. Only prednisone is able to restore abnormally high stretch closer to wild-type levels; serotonin and melatonin have no significant effect. These results in the swim environment partially mirror the result of pharmacological treatments in improving strength: although the compounds do have advantageous effects on deficiencies, treated *dys-1* animals still trail behind wild-type animals. Furthermore, the resistance of animal asymmetry to the various treatments and the resistance of stretch to two of the three treatments indicate that these compounds are not targeting all aspects of physiological abnormalities in the *dys-1* mutants.

**Fig. 5.**
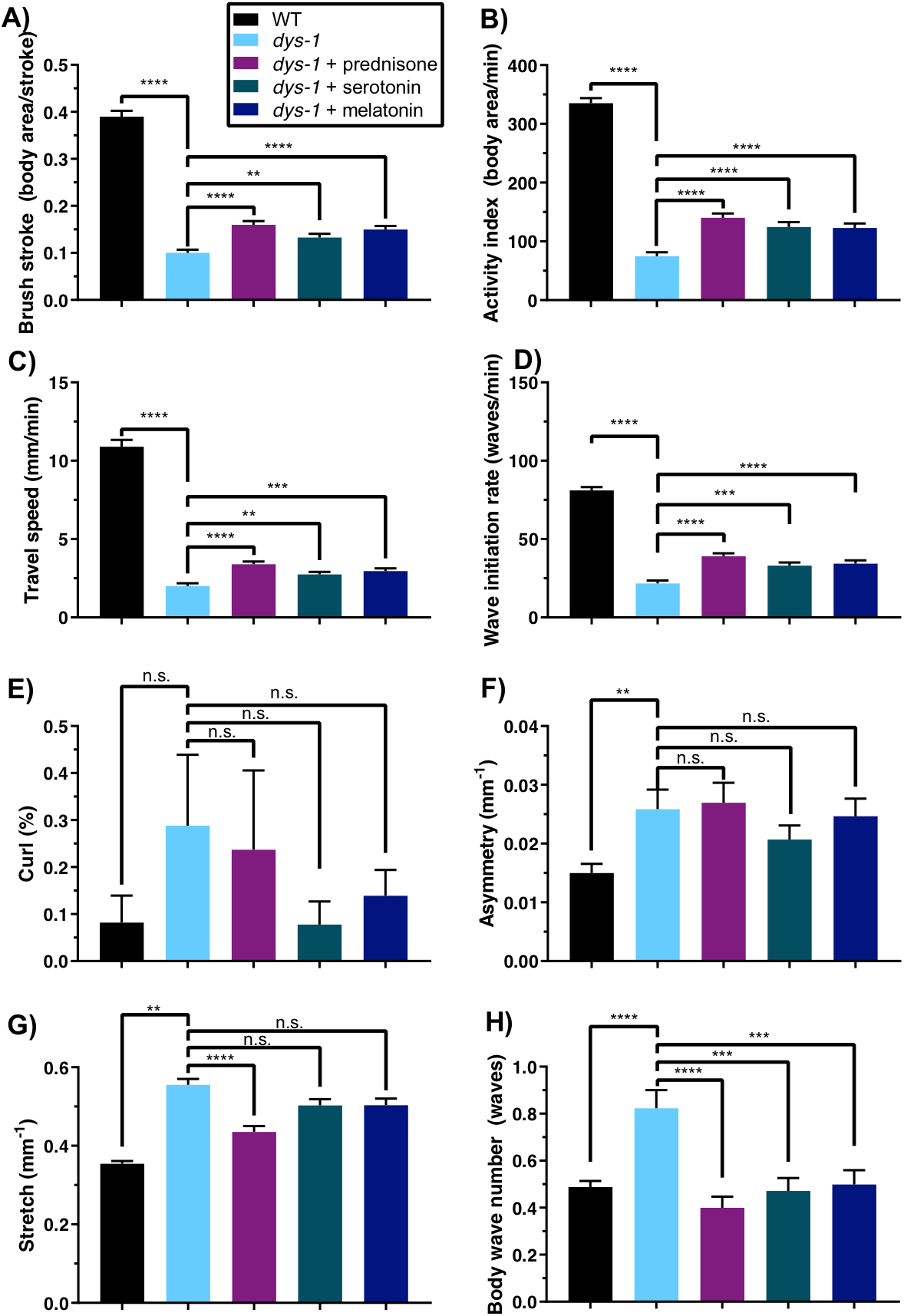
Swimming environment: *dys-1(eg33)* mutants are deficient in nearly all swimming measures, and treatments with prednisone, serotonin, and melatonin improve most measures. *dys-1(eg33)* mutants have significantly lower (A) brush stroke, (B) activity index, (C) travel speed, and (D) wave initiation rate compared to wild type. Treatments with prednisone, serotonin, and melatonin give significant improvements in these measures. (E) *dys-1(eg33)* mutants are not significantly different from wild type in their curling percentage, but their (F) asymmetry, (G) stretch, and (H) body wave number are significantly higher than those of wild type, and all compounds improve these measures, with the exception of asymmetry, which is not improved under any treatment, and stretch, which is improved only by prednisone. All four of these measures, which reflect body posture, increase with age in wild-type animals. WT: N=56; *dys-1*: N=56; *dys-1*+prednisone: N=55; *dys-1*+serotonin: N=54; *dys-1*+melatonin: N=56. Significance between WT and *dys-1* controls was assessed with a 2-sample t-test; significance between *dys-1* and *dys-1*+treatments was assessed with one-way ANOVA with Dunnett’s post-hoc test. n.s., not significant; **, P≤0.01; ***, P≤0.001; ****, P≤0.0001.

#### Burrowing environment

*dys-1(eg33)* mutants exhibit burrowing defects [31, 43], which we sought to confirm using our novel hydrogel-based burrowing platform. Here, we assessed the burrowing ability of *dys-1* animals that had been treated with pharmacological interventions, which is the first report of how pharmacological interventions impact burrowing ability in *C. elegans*. We find that while over 80% of wild-type animals reach the surface by the end of the 2-hour burrowing assay, only about 5% of *dys-1* mutants reached the surface during this same time frame (Figure 6). The strong defect in *dys-1* burrowing ability suggests that burrowing may place a high demand on the muscle and requires intact excitation-contraction coupling in the muscle. These defects are only partially addressed by prednisone, melatonin, and serotonin, as each compound gives significant but modest improvements in the burrowing ability of *dys-1*, although treated animals still fall short of wild-type performance (Figure 6). Our data support the notion that the burrowing assay enhances the dynamic range in which we can evaluate drug interventions that counter *dys-1* deficits.

**Fig. 6.**
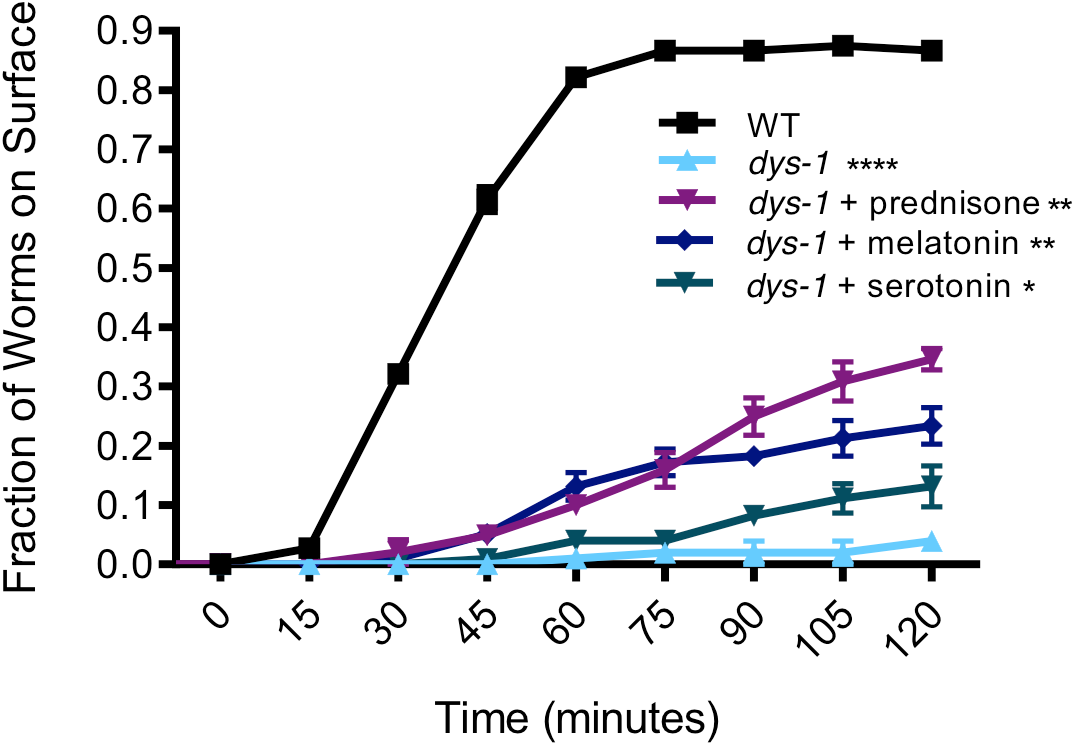
Burrowing environment: dystrophin mutants are highly deficient in burrowing and some interventions modestly improve this deficiency. *dys-1(eg33)* mutants are highly deficient in burrowing, with only less than 10% of the population able to reach the surface compared to over ~80% of wild-type animals after a 2 hour trial. Prednisone, melatonin, and serotonin confer a significant improvement in the burrowing ability of *dys-1(eg33)* mutants, although their performance still falls short of that of wild type. Trials were in triplicate with average samples sizes of: WT: N=37; dys-1: N=34; dys-1+prednisone: N=34; dys-1+serotonin: N=33; dys-1+melatonin: N=33. Significance was assessed with two-way ANOVA. n.s., not significant; *, P≤0.05; **, P≤0.01; ****, P≤0.0001.

## DISCUSSION

### A novel multi-environment phenotyping platform

Assessment of physical performance in humans is not only central to a variety of disease contexts but is also critical for evaluating the effects of factors such as aging, diet, and exercise. Development of measures that are indicative of physical performance in *C. elegans* that are analogous to human measures is expected to increase the translational relevance of this genetic model. Behavioral assays that extract descriptive measures of *C. elegans* swim locomotion [26, 35, 36], crawling on NGM plates [22, 27, 44–46], and burrowing [24, 31] have been documented, but none have investigated individuals in these different environments in the context of assessing physical performance. We suggest that our approach that integrates multiple measures from three unique physical environments advances the general field of phenotyping in *C. elegans*.

In this study, we establish the value of this MEP platform by showing that the parameters reported are not redundant with one another—testing in multiple environments increases the “bandwidth” over which deficits can be detected and improvements can be documented. Our gene expression data show that swimming, crawling, and burrowing place unique demands on the nematode, as evidenced by analysis of a sample set of genes representative of oxidative stress response, glucose metabolism, and fat metabolism, which are differentially expressed depending on the animal’s environment and specific locomotory response. The environments of swimming, crawling, and burrowing can put different demands on the organism (e.g. oxygen, pressure, and physical forces) and its physiology (muscle demands and type of movement).

### Application of MEP to *C. elegans* DMD model

Previous work reports studies of *dys-1* mutant deficits using single parameters such as crawling speeds [47], swimming speeds [48], burrowing ability [31], or muscle strength [37]. Additional work that did give multiple readouts of *dys-1* or *dys-1;hlh-1* animal physiology established baseline values for a few additional parameters within the swimming environment using the Biomechanical Profiling (BMP) platform but did not look at the effects of any interventions [26, 49]. More recently, one study evaluated *dys-1* mutants in swimming, crawling, and burrowing environments and focused solely on the frequencies and amplitudes of the animal motion in the different environments [43].

Building on this prior work, we applied the MEP platform to test whether additional deficiencies could be detected in the same animal populations when multiple phenotypic measures were assayed. While crawling activity, muscular endurance, and curl percentage are not significantly different in *dys-1* mutants as compared with their wild-type counterparts, all other nine measures we report are strikingly worse. These nine measures span each of the three physical environments, indicating that we can detect deficiencies in the same animal population in all environments comprising the MEP platform. To establish the efficacy of the platform in detecting improvements from pharmacological interventions, we assessed animals that had been treated with prednisone and melatonin, previously reported to improve muscle strength [37], as well as serotonin, which had been previously reported to reduce the number of damaged muscle cells in the *dys-1;hlh-1* DMD model [41, 50].

The findings from our study here inform two potential directions in researching therapeutic compounds for DMD. First, any compounds that counter defects in all the measures reported in the MEP platform would be most promising to pursue in *mdx* mice and eventually clinical trials. Second, compounds that restore a subset of the deficiencies to wild-type levels may be worth pursuing and might be easily optimized in the facile *C. elegans* model. Similar to our study with the DMD model, future opportunities also exist to test other *C. elegans* disease models in which the neuromuscular system is impaired to potentially identify candidates for therapeutic intervention.

## CONCLUSIONS

In this work, we have shown the power of a multi-environmental phenotyping platform that holds potential for improving translation of findings in *C. elegans* disease models to humans. The three different physical environments, which elicit distinct locomotory modes of crawling, swimming, and burrowing, each also challenge the animal in distinct manners, as demonstrated by the unique information extracted from each environment and the differences in expression of select genes in each environment. We demonstrated the utility of this framework by assessing the health of the *C. elegans* model for Duchenne muscular dystrophy, both with and without pharmacological interventions. The framework we describe here can be utilized in the future for robustly assessing health and determining interventions that might be most successfully translated from *C. elegans* disease models to mammalian systems.

## MATERIALS AND METHODS

### Nematode strains, culture, and drug treatments

Strains used in this experiment were wild-type (N2 strain) and *dys-1(eg33)* (BZ33 strain). The strains were provided by the *Caenorhabditis* Genetics Center (CGC). For all drug treatments, stock solutions were aliquoted on the surface of seeded NGM plates approximately 24 hours before using the plates. The final concentration of the drug was calculated based on the total volume of the contents of the plate. Concentrations used for drug treatments were matched with concentrations used in previous studies: 0.37 mM prednisone [37], 1 mM melatonin [37], 2.5 mM serotonin creatinine sulfate [41]. A control solvent was added to the surface of agar plates used for culturing animals that did not receive a drug treatment.

Synchronized *C. elegans* populations were prepared by picking ~20-30 adult animals to each seeded plate and allowing them to lay eggs for ~3-4 hours. At the end of this time period, adult animals were picked and discarded, leaving synchronized eggs on the plate. Eggs were allowed to hatch and grow at 20°C until the first or second day of adulthood, when all assays were conducted. Animals for all assays were Day 2 adults, except for animals used for gene expression and burrowing experiments, which were Day 1 adults. For assays done on Day 2 of adulthood, animals were transferred on Day 1 to fresh plates in order to ensure sufficient food and minimal crowding.

### NemaFlex strength assay

The NemaFlex strength assay was conducted as previously described, with a few modifications [34, 37]. In our previous study, we used a microfluidic device design that was optimized to accommodate animals of a much wider age range [37]. For the present study, we instead utilized a device with modified pillar diameter and spacing that is best used for Day 1 and Day 2 adults. Here, the pillars had a smaller diameter (~41μm) and tighter spacing (74 μm between edges of the pillars) due to the smaller size of the worms at the beginning of adulthood. Because of this, the force constants calculated from this device are not as large as those calculated using the device in our previous study [37]; this therefore results in smaller force values. However, the same device is used for all experiments within this study, which allows for consistency in the comparisons of the data presented here.

Synchronized animals were loaded into the devices with one animal per chamber. A 60-second video was collected for each animal using a Nikon TI-E microscope with a 4x objective and Andor Zyla sCMOS 5.5 camera at a frame rate of 5 FPS. Movies were then processed and analyzed for force values using our in-house-built image processing software (MATLAB, R2015b). Animal strength is determined by selecting for the maximum force exerted in each frame (300 total frames and therefore force values per animal), and then selecting for the 95^th^ percentile of these forces. The 95^th^ percentile is selected rather than the absolute maximum, as it is more robust to error. Two new measures presented here are endurance and crawling activity. Endurance is calculated in a similar way to strength, but instead the 50^th^ percentile of the maximum forces exerted is normalized by the strength, or the 95^th^ percentile. Thus, this is the average force exerted normalized by our definition of strength and gives an idea of the profile of forces being exerted by the animal. Crawling activity is defined as the number of pillars that the animal interacts with per second, where pillars are automatically counted by image processing software, and the count of the pillars only needs to be normalized to the duration of the acquired movie. Significances between wild type and *dys-1* controls were assessed using a 2-sample *t*-test. Significance between *dys-1* and *dys-1*+treatments was assessed with one-way ANOVA with Dunnett’s post-hoc test.

### CeleST swim assay

*<u>C.</u> <u>ele</u>gans* <u>S</u>wim <u>T</u>est (CeleST) assays were conducted as previously described [35, 36]. Animals were picked and placed into a 50 μL aliquot of M9 Buffer on a specialized glass slide with two, 10-mm pre-printed rings on the surface (Thermo Fisher Scientific), as previously used by the original developers of the CeleST assay. The rings hold in the M9 buffer and act as a swim arena for the animals. Four to five animals were placed into each M9 buffer aliquot, and two swim arenas were loaded at a time and then imaged sequentially. Images were acquired with a Nikon TI-E microscope and Andor Zyla sCMOS 5.5 camera at a frame rate of ~12 FPS. Images were acquired using dark-field imaging to allow a bright worm on a dark background. Images were processed using the previously developed CeleST software, although the most recent version of the software includes 8 of the originally reported 10 measures; the measures of reverse swimming and attenuation have been dropped from recent releases of the CeleST software and were thus not considered for the experiments described here. The CeleST software reports the percentage of frames where a worm was successfully and accurately segmented, and only these valid frames are used towards the calculation of all measures. An approximate validity cutoff of 80% was implemented to ensure that sufficient and continuous frames were available for measure computation. Significances between wild type and *dys-1* controls were assessed using a 2-sample *t*-test. Significance between *dys-1* and *dys-1*+treatments was assessed with one-way ANOVA with Dunnett’s post-hoc test.

### Pluronic gel-based burrowing assay

Burrowing assays were conducted as previously described [25]. Briefly, approximately 30 to 40 *C. elegans* were introduced into the base of a well of a 12-well plate by handpicking with a platinum wire into a small pluronic drop. Transfer of *E. coli* to the well plate was avoided by first transferring animals to an unseeded NGM plate and letting them crawl for a few minutes before the animals were added to the well plate. A volume of 2.5 mL of pluronic was added to form the upper layer of the gel, which gave a gel height of about 0.75 cm. The exact number of animals loaded into the well was then counted and recorded. Once the gel had solidified, 20 μL concentrated *E. coli* solution was added to the surface above the animals. The concentrated *E. coli* solution was made by re-suspending an *E. coli* pellet in liquid NGM, which is just NGM without the agar. The addition of the *E. coli* solution marked the *t = 0* time point of the assay. The number of animals on the surface was then scored every 15 minutes for a total of 2 hours unless otherwise noted. The percent of the population reaching the surface at each time point was then calculated. For all experiments conducted, a concentration of 26 w/w% of Pluronic F-127 (Sigma-Aldrich) in DI water was used. Each condition/strain was run in triplicate, and significances were assessed using 2-way ANOVA in GraphPad Prism software.

### Individual-level, multi-environment phenotyping

NemaFlex, CeleST, and burrowing assays were conducted as previously described but with a few modifications that enabled sustained tracking of the individual animal identity, which allowed looking at each individual animal under burrowing, crawling, and swimming environments. Animals were loaded into the 12-well plates as described, and the addition of bacteria marked the start of the assay. Rather than counting the number of animals on the surface, however, individual animals were collected from the burrowing wells as they reached the surface. Each animal was given a score of how many minutes it took to reach the surface. Individual animals were placed in a 35mm NGM plate with *E. coli* OP50 and allowed to recover for 4 hours. Individual animals were then imaged swimming for CeleST analysis (swimming only for a few minutes), and then were placed back on their agar plate. After all imaging was finished for swimming animals, individual worms were loaded into NemaFlex chambers and imaged and analyzed for strength, activity, and muscular endurance as previously described. The only difference in data representation is for that of burrowing ability, where individual animals were given the score of number of minutes taken to reach the surface. This is in contrast to the typical representation of the percent of the population that has reached the surface at a given time point that is reported when populations rather than individuals are assessed.

### Gene expression via quantitative PCR

To assess how *C. elegans* respond to the three different environments of crawling, burrowing, and swimming, we conducted an assay where wild-type *C. elegans* were allowed to burrow for 90 minutes, and then we assessed the genetic response as related to oxidative stress, glucose metabolism, and fat metabolism. Animals were loaded into burrowing wells as previously described, except a tall gel layer that filled the well completely to the top was added to ensure that animals did not exit the burrowing environment during the 90 minutes of burrowing. Liquid NGM was added to the surface to provide chemotactic stimulation to the animals without introducing a food source (*E. coli)*. A control group of animals were transferred to unseeded NGM plates for the 90-minute duration of the burrowing. At the conclusion of the 90 minutes, approximately 30 each of burrowing and control animals were collected in M9 Buffer. Another set of burrowing and control animals were transferred to seeded NGM plates for recovery, and then were collected after 2 hours.

The RNA extraction and qPCR protocols were consistent with those previously described [38, 51]. Immediately after collection in M9 Buffer, animals were spun down and the supernatant was removed. After the addition of TRIzol (Ambion), animals underwent snap freezing with liquid nitrogen. The samples then underwent freeze/thaw cycles with liquid nitrogen and a heat block at 37°C. Subsequently, total RNA was extracted as instructed by the manufacturer’s protocol (Ambion), and cDNA was made with the SuperScript III First-Strand Synthesis System (Invitrogen). The PerfeCTa SYBR Green FastMix (Quantabio), 0.5 μM of primers for each gene, and diluted cDNA were used for conducting qPCR. The primers used match those previously published as part of the prior analysis done for gene expression changes after 90 minutes of swimming [38]. A 7500 Fast Real-Time PCR System (Applied Biosystems) was used with the ΔΔCt method relative expression method for calculations [52]. For reference genes, *cdc-42* and Y45F10D.4 were used [53].

#### Statistical analyses

All significances were assessed with a 2-sample t-test, except for population burrowing comparisons, which were assessed using two-way ANOVA in GraphPad Prism software, and NemaFlex and CeleST measures for *dys-1* vs. *dys-1*+treatments, which were assessed with one-way ANOVA with Dunnett’s post-hoc test.

## ACKNOWLEDGEMENTS

Some strains were provided by the CGC, which is funded by NIH Office of Research Infrastructure Programs (P40 OD010440). We would like to acknowledge Leila Lesanpezeshki for useful discussions surrounding burrowing experiments and for fabricating the mold used to make NemaFlex microfluidic devices. This work is partially supported by funding from the National Institutes of Health (RO1 AG051995-04 to M.D. & S.V.), National Aeronautics and Space Administration (NNX15AL16G to S.V.), and the Biotechnology and Biological Sciences Research Council (BB/N015894/1 to N.J.S.). J.E.H. acknowledges funding support from the Fulbright U.S. Student Program and the Germanistic Society of America. R.L. has been funded by postdoctoral fellowships from Life Sciences Research Foundation (sponsored by Simons Foundation; award # Laranjeiro-2015) and American Heart Association (award # 18POST33960502). A.A. was supported by the Max Planck Society.

## REFERENCES

1. Caspersen CJ, Powell KE, Christenson GM. Physical activity, exercise, and physical fitness: definitions and distinctions for health-related research. Public health rep. 1985;100(2):126–31.

2. McDonald CM. Physical activity, health impairments, and disability in neuromuscular disease. American journal of physical medicine & rehabilitation. 2002;81(11):S108–S20.

3. Beaudart C, Rolland Y, Cruz-Jentoft AJ, Bauer JM, Sieber C, Cooper C, et al. Assessment of muscle function and physical performance in daily clinical practice. Calcified Tissue International. 2019:1–14.

4. Wang L, Larson EB, Bowen JD, van Belle G. Performance-based physical function and future dementia in older people. Archives of internal medicine. 2006;166(10):1115–20.

5. Guralnik JM, Simonsick EM, Ferrucci L, Glynn RJ, Berkman LF, Blazer DG, et al. A short physical performance battery assessing lower extremity function: association with self-reported disability and prediction of mortality and nursing home admission. Journal of gerontology. 1994;49(2):M85–M94.

6. Studenski S, Perera S, Wallace D, Chandler JM, Duncan PW, Rooney E, et al. Physical performance measures in the clinical setting. Journal of the American Geriatrics Society. 2003;51(3):314–22.

7. Volpato S, Cavalieri M, Sioulis F, Guerra G, Maraldi C, Zuliani G, et al. Predictive value of the Short Physical Performance Battery following hospitalization in older patients. The Journals of Gerontology Series A: Biological Sciences and Medical Sciences. 2011;66(1):89–96.

8. Rockwood K, Song X, MacKnight C, Bergman H, Hogan DB, McDowell I, et al. A global clinical measure of fitness and frailty in elderly people. Canadian Medical Association Journal. 2005;173(5):489–95.

9. Brenner S. The genetics of *Caenorhabditis elegans*. Genetics. 1974;77(1):71–94.

10. Collins CA, Morgan JE. Duchenne’s muscular dystrophy: animal models used to investigate pathogenesis and develop therapeutic strategies. International Journal of Experimental Pathology. 2003;84(4):165–72.

11. Nass R, Miller DM, Blakely RD. *C. elegans*: a novel pharmacogenetic model to study Parkinson’s disease. Parkinsonism & Related Disorders. 2001;7(3):185–91.

12. Harrington AJ, Hamamichi S, Caldwell GA, Caldwell KA. *C. elegans* as a model organism to investigate molecular pathways involved with Parkinson’s disease. Developmental Dynamics. 2010;239(5):1282–95.

13. Link CD. *C. elegans* models of age-associated neurodegenerative diseases: lessons from transgenic worm models of Alzheimer’s disease. Experimental Gerontology. 2006;41(10):1007–13.

14. Parker JA, Connolly JB, Wellington C, Hayden M, Dausset J, Neri C. Expanded polyglutamines in *Caenorhabditis elegans* cause axonal abnormalities and severe dysfunction of PLM mechanosensory neurons without cell death. Proceedings of the National Academy of Sciences. 2001;98(23):13318–23.

15. Poulin G, Nandakumar R, Ahringer J. Genome-wide RNAi screens in *Caenorhabditis elegans:* impact on cancer research. Oncogene. 2004;23(51):8340–5.

16. Herndon LA, Schmeissner PJ, Dudaronek JM, Brown PA, Listner KM, Sakano Y, et al. Stochastic and genetic factors influence tissue-specific decline in ageing *C. elegans*. Nature. 2002;419(6909):808–14.

17. Bushby K, Finkel R, Birnkrant DJ, Case LE, Clemens PR, Cripe L, et al. Diagnosis and management of Duchenne muscular dystrophy, part 1: diagnosis, and pharmacological and psychosocial management. The Lancet Neurology. 2010;9(1):77–93.

18. Folch J, Petrov D, Ettcheto M, Abad S, Sánchez-López E, García ML, et al. Current research therapeutic strategies for Alzheimer’s disease treatment. Neural Plasticity. 2016;2016.

19. Dehay B, Bourdenx M, Gorry P, Przedborski S, Vila M, Hunot S, et al. Targeting α-synuclein for treatment of Parkinson’s disease: mechanistic and therapeutic considerations. The Lancet Neurology. 2015;14(8):855–66.

20. Ross CA, Tabrizi SJ. Huntington’s disease: from molecular pathogenesis to clinical treatment. The Lancet Neurology. 2011;10(1):83–98.

21. Kwak JY, Kwon K-S. Pharmacological Interventions for Treatment of Sarcopenia: Current Status of Drug Development for Sarcopenia. Journal of the Korean Geriatrics Society. 2019.

22. Nussbaum-Krammer CI, Neto MF, Brielmann RM, Pedersen JS, Morimoto RI. Investigating the spreading and toxicity of prion-like proteins using the metazoan model organism *C. elegans*. JoVE (Journal of Visualized Experiments). 2015;(95):e52321.

23. Buckingham SD, Sattelle DB. Fast, automated measurement of nematode swimming (thrashing) without morphometry. Bmc Neuroscience. 2009;10(1):1.

24. Bainbridge C, Schuler A, Vidal-Gadea AG. Method for the assessment of neuromuscular integrity and burrowing choice in vermiform animals. Journal of Neuroscience Methods. 2016;264:40–6.

25. Lesanpezeshki L, Hewitt JE, Laranjeiro R, Antebi A, Driscoll M, Szewczyk NJ, et al. Pluronic gel-based burrowing assay for rapid assessment of neuromuscular health in *C. elegans*. Scientific Reports. 2019;(1):15246. doi: 10.1038/s41598-019-51608-9.

26. Krajacic P, Shen X, Purohit PK, Arratia P, Lamitina T. Biomechanical profiling of *Caenorhabditis elegans* motility. Genetics. 2012;191(3):1015–21.

27. Yemini E, Jucikas T, Grundy LJ, Brown AEX, Schafer WR. A database of *Caenorhabditis elegans* behavioral phenotypes. Nature Methods. 2013;10(9):877–9.

28. Fang-Yen C, Wyart M, Xie J, Kawai R, Kodger T, Chen S, et al. Biomechanical analysis of gait adaptation in the nematode Caenorhabditis elegans. Proceedings of the National Academy of Sciences. 2010;107(47):20323–8.

29. Shen XN, Sznitman J, Krajacic P, Lamitina T, Arratia PE. Undulatory locomotion of Caenorhabditis elegans on wet surfaces. Biophysical journal. 2012;102(12):2772–81.

30. Bilbao A, Patel AK, Rahman M, Vanapalli SA, Blawzdziewicz J. Roll maneuvers are essential for active reorientation of Caenorhabditis elegans in 3D media. Proceedings of the National Academy of Sciences. 2018:201706754.

31. Beron C, Vidal-Gadea AG, Cohn J, Parikh A, Hwang G, Pierce-Shimomura JT. The burrowing behavior of the nematode *Caenorhabditis elegans*: a new assay for the study of neuromuscular disorders. Genes, Brain and Behavior. 2015;14(4):357–68.

32. Johari S, Nock V, Alkaisi MM, Wang W. On-chip analysis of *C. elegans* muscular forces and locomotion patterns in microstructured environments. Lab on a Chip. 2013;13(9):1699–707.

33. Khare SM, Awasthi A, Venkataraman V, Koushika SP. Colored polydimethylsiloxane micropillar arrays for high throughput measurements of forces applied by genetic model organisms. Biomicrofluidics. 2015;9(1):014111.

34. Rahman M, Hewitt J, Van-Bussel F, Edwards H, Blawzdziewicz J, Szewczyk N, et al. NemaFlex: A microfluidics-based technology for standardized measurement of muscular strength of *C. elegans*. Lab on a Chip. 2018;18(15):2187–201. doi: 10.1039/C8LC00103K.

35. Restif C, Ibáñez-Ventoso C, Vora MM, Guo S, Metaxas D, Driscoll M. CeleST: computer vision software for quantitative analysis of *C. elegans* swim behavior reveals novel features of locomotion. PLOS Computational Biology. 2014.

36. Ibáñez-Ventoso C, Herrera C, Chen E, Motto D, Driscoll M. Automated Analysis of *C. elegans* Swim Behavior Using CeleST Software. JoVE (Journal of Visualized Experiments). 2016;(118):e54359–e.

37. Hewitt JE, Pollard AK, Lesanpezeshki L, Deane CS, Gaffney CJ, Etheridge T, et al. Muscle strength deficiency and mitochondrial dysfunction in a muscular dystrophy model of *Caenorhabditis elegans* and its functional response to drugs. Disease Models & Mechanisms. 2018;11(12):dmm036137.

38. Laranjeiro R, Harinath G, Burke D, Braeckman BP, Driscoll M. Single swim sessions in *C. elegans* induce key features of mammalian exercise. BMC Biology. 2017;15(1):30. doi: 10.1186/s12915-017-0368-4.

39. DeSilva S, Drachman DB, Mellits D, Kuncl RW. Prednisone treatment in Duchenne muscular dystrophy: long-term benefit. Archives of Neurology. 1987;44(8):818–22.

40. Chahbouni M, Escames G, Venegas C, Sevilla B, García JA, Lopez LC, et al. Melatonin treatment normalizes plasma pro-inflammatory cytokines and nitrosative/oxidative stress in patients suffering from Duchenne muscular dystrophy. Journal of Pineal Research. 2010;48(3):282–9.

41. Carre-Pierrat M, Mariol M-C, Chambonnier L, Laugraud A, Heskia F, Giacomotto J, et al. Blocking of striated muscle degeneration by serotonin in *C. elegans*. Journal of Muscle Research & Cell Motility. 2006;27(3-4):253–8.

42. Gurel V, Lins J, Lambert K, Lazauski J, Spaulding J, McMichael J. Serotonin and Histamine Therapy Increases Tetanic Forces of Myoblasts, Reduces Muscle Injury, and Improves Grip Strength Performance of Dmdmdx Mice. Dose-Response. 2015;13(4):1559325815616351.

43. Hughes KJ, Rodriguez A, Flatt KM, Ray S, Schuler A, Rodemoyer B, et al. Physical exertion exacerbates decline in the musculature of an animal model of Duchenne muscular dystrophy. Proceedings of the National Academy of Sciences. 2019;116(9):3508–17.

44. Ghosh R, Mohammadi A, Kruglyak L, Ryu WS. Multiparameter behavioral profiling reveals distinct thermal response regimes in *Caenorhabditis elegans*. BMC Biology. 2012;10(1):85.

45. Feng Z, Cronin CJ, Wittig JH, Sternberg PW, Schafer WR. An imaging system for standardized quantitative analysis of *C. elegans* behavior. BMC Bioinformatics. 2004;5(1):115.

46. Nahabedian JF, Qadota H, Stirman JN, Lu H, Benian GM. Bending amplitude–A new quantitative assay of *C. elegans* locomotion: Identification of phenotypes for mutants in genes encoding muscle focal adhesion components. Methods. 2012;56(1):95–102.

47. Oh KH, Kim H. Reduced IGF signaling prevents muscle cell death in a *Caenorhabditis elegans* model of muscular dystrophy. Proceedings of the National Academy of Sciences. 2013;110(47):19024–9.

48. Yuan J, Ko H, Raizen DM, Bau HH. Terrain following and applications: *Caenorhabditis elegans* swims along the floor using a bump and undulate strategy. Journal of The Royal Society Interface. 2016;13(124):20160612.

49. Sznitman J, Purohit PK, Krajacic P, Lamitina T, Arratia PE. Material properties of *Caenorhabditis elegans* swimming at low Reynolds number. Biophysical Journal. 2010;98(4):617–26.

50. Gaud A, Simon J-M, Witzel T, Carre-Pierrat M, Wermuth CG, Ségalat L. Prednisone reduces muscle degeneration in dystrophin-deficient *Caenorhabditis elegans*. Neuromuscular Disorders. 2004;14(6):365–70.

51. Laranjeiro R, Harinath G, Hewitt JE, Hartman JH, Royal MA, Meyer JN, et al. Swim exercise in Caenorhabditis elegans extends neuromuscular and gut healthspan, enhances learning ability, and protects against neurodegeneration. Proceedings of the National Academy of Sciences. 2019;116(47):23829–39.

52. Livak KJ, Schmittgen TD. Analysis of relative gene expression data using real-time quantitative PCR and the 2− ΔΔCT method. Methods. 2001;25(4):402–8.

53. Hoogewijs D, Houthoofd K, Matthijssens F, Vandesompele J, Vanfleteren JR. Selection and validation of a set of reliable reference genes for quantitative sod gene expression analysis in *C. elegans*. BMC Molecular Biology. 2008;9(1):9.

